# Demographic insights for coral restoration

**DOI:** 10.1101/2025.10.21.683544

**Authors:** Joshua S. Madin, Thomas Oliver, Mike McWilliam, Mollie Asbury, Andrew H. Baird, Guanyan Keelung Chen, Sean R. Connolly, Courtney Couch, Crawford Drury, Jon Ehrenberg, Hendrikje Jorissen, Allison D. Nims, Jessica Reichert, Lomani Rova, Nina M. D. Schiettekatte, Robert J. Toonen, Devynn M. Wulstein

## Abstract

1. Coral reef decline has prompted a global surge in reef restoration initiatives. The success of initiatives that aim to sustain coral populations or assemblages will depend on demographic principles. Restoration strategies generally follow two demographic pathways: supplemental approaches, which increase population numbers through the repeated addition of recruits, fragments, or adults, without fundamentally altering long-term population dynamics; and structural approaches, which enhance vital rates–growth, survival, or reproduction– through changes that modify the demographic processes governing population growth on a sustained basis, either extrinsically (e.g., herbivory, habitat protection) or intrinsically (e.g., assisted evolution). Some interventions, such as supplemental feeding, may temporarily improve vital rates but still function as supplemental approaches because benefits persist only while interventions continue.
2. We synthesized 28 coral matrix population models spanning morphologically diverse coral species from the Caribbean, Hawaii, and the Great Barrier Reef to quantify supplemental and structural changes needed to increase population growth.
3. Results highlight two consistent demographic leverage points for which population growth was most sensitive: (1) survival of reproductive adults and (2) successful recruitment. Improving adult survival or recruitment by 20% through structural means produced a 5% increase in population growth rate across population on average, assuming the whole population was affected. By contrast through supplemental means, the same increase required ∼100 recruit outplants or 5–10 adult outplants per 1,000 individuals in a population annually. For large populations typical of restoration targets (10^5^–10^7^ individuals), this translates to 10^3^ adult and 10^4^ recruit outplants per year—levels rarely logistically or economically feasible.
4. These findings yield two key implications. First, supplemental interventions are inefficient for large populations and demand sustained, large-scale effort, even when they temporarily enhance growth or survival. Second, strategies enhancing vital rates across broad geographic areas represent the most effective means of boosting coral abundance, including habitat protection, alleviation of environmental stressors, and interventions which promote long-term survival and recruitment. Our demographic framework underscores that if the restoration goal is sustained increases in coral cover, success depends less on repeated supplements and more on interventions that produce lasting improvements in vital rates.

## Introduction

Coral reef conservation is transitioning from a focus on species and habitat protection to active restoration (Rinkevich 2019). Many scientists are skeptical that coral restoration can change the course of the coral reef crisis (Hughes et al. 2023), in part because of the vast geographical areas of reefs, numerous coral species, diverse life history strategies, and large coral population sizes (McWilliam et al. 2018, Dietzel et al. 2021). However, modeling efforts suggest that some local-scale actions combined with large-scale action on climate change could result in significantly different reef outcomes (Bay et al. 2017, Committee on Interventions to Increase the Resilience of Coral Reefs 2019, McManus et al. 2021). Indeed, the numbers and scales of coral reef restoration initiatives are growing rapidly (e.g., SECORE, Coral Nurture Program, RRAP, MARS, reviewed by Boström-Einarsson et al. 2020; McLeod at al. 2022; Edwards et al. 2024). Governments and nonprofits are spending millions of dollars on research and logistics supporting restoration (Bayraktarov et al. 2019, Tortolero-Langarica et al. 2020; Sam et al. 2021, Dela Cruz & Harrison 2017; reviewed by Watt-Pringle et al. 2024; Segaran et al. 2024; Schmidt-Roach et al. 2025). Assessments of restoration practices are generating mixed reviews among ecologists (Boström-Einarsson et al. 2020, McLeod et al. 2022; Madin et al. 2023, Hughes et al. 2023, Streit et al. 2024; Suggett et al. 2024; Watt-Pringle et al. 2024). Meanwhile, corals continue to fare poorly (Hoegh-Guldberg et al. 2023).

Coral population growth or decline can be projected quantitatively as a matrix of transitions among categorical life history stages (Hughes 1984), or through a continuous range of sizes using an Integral Projection Model (e.g., McWilliam et al. 2023). Like most plants and animals, colonial reef building corals are often classified simply into three life history stages (Caswell 1989). The first is the recruitment stage, which represents new individuals entering the population by sexual or asexual reproduction (Hughes and Jackson 1980). Recruits tend to have low growth rates (absolute change in colony area) and exceptionally low survival rates (Carlot et al. 2021). Second is the juvenile stage, which is generally considered the point prior to reproductive maturity at which a colony is large enough to reduce post-settlement mortality and growth becomes steady and predictable (Stearns 1989, Doropoulos et al. 2016). Third is the adult stage where growth slows and colonies become reproductively mature. Reproductive output scales with the number of polyps which, in turn, scales with colony size (Hall and Hughes 1996; Padilla-Gamiño & Gates 2012; Álvarez-Noriega et al 2016). A complete population matrix describes transitions between each of these stages (including recruitment, growth, shrinkage, and survival, Fig. 1A), and the dominant eigenvalue of the matrix (known as lambda, λ) reflects the intrinsic growth rate of the population, which can be used as a common currency for comparing population dynamics (Edmunds et al. 2014; Cant et al. 2021). Values above one (λ > 1) translate to population growth, and those below one (λ < 1) indicate population decline. Lambda is also a metric of population resilience following disturbances, as it reflects the rate at which a population recovers to pre-disturbance levels (Madin et al. 2012; McWilliam et al. 2023). Therefore, it can be a useful tool for guiding conservation decisions (Crouse et al. 1987; Doak et al. 2021; Edmunds et al. 2014; Edmunds & Riegl 2020).

**Figure 1.**
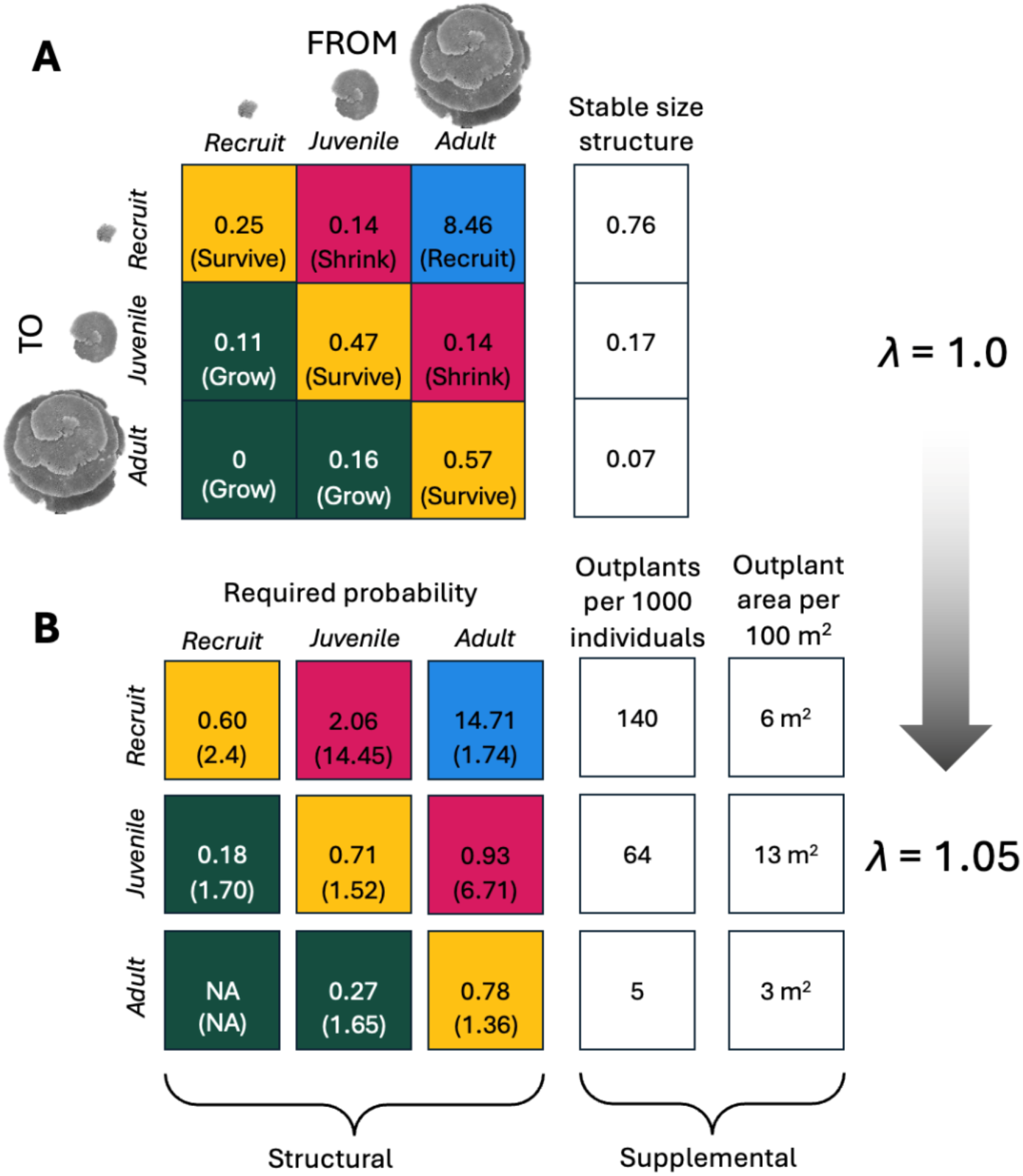
The yearly stage transitions for a population of *Acropora cytherea* (sensu Veron 2000) at Lizard Island on the Great Barrier Reef. (**A**) Most cells show probabilities of moving between stages in one year. The blue cell represents the number of recruits that return to the population per year from a single adult for the population growth to be at stasis (λ=1.0). The stable size structure is the distribution of proportions of colonies in each stage as the population approaches its long-run growth rate. (**B**) The transition cell value required to increase the intrinsic population growth rate to λ=1.05, a 5% increase. (Proportional changes are given in parentheses.) Note that cell values greater than 1 are not possible, except for the blue cell. On the right are the number of individuals and outplant cover required for each stage to increase λ=1.05 without changing transition probabilities.

We consider management interventions broadly as falling on a continuum between what we term “structural” and “supplemental” interventions. Under traditional coral reef management, such interventions have been conducted by improving the environment (e.g., reducing pollution [Vega Thurber et al. 2014; Lapointe et al. 2019; Gove et al. 2023], and removing competitors [Ceccarelli et al. 2018] and predators [Pratchett et al. 2017, Raker et al. 2023]). Nevertheless, as a response to the increasingly negative effects of climate change, the focus has shifted to more active interventions such as moving corals or gametes between environments (assisted migration [Vitt et al. 2009; Hagedorn et al. 2021; Barreto et al. 2023]) and selective breeding (assisted evolution [van Oppen et al. 2015; Humanes et al. 2021; Drury et al. 2022]), or by supplementing population numbers via outplanting of coral larvae, fragments, or whole colonies (Boström-Einarsson et al. 2020; Edwards et al. 2024).

Within this diverse set of intervention approaches, we consider an intervention “structural” when an intervention over a defined period produces changes to a population’s vital rates that are sustained over many generations beyond the time frame of the active intervention. Thus, for example, deployment of artificial habitat that does not require ongoing maintenance increases available settlement space for new recruits for the life of the artificial structure (presumably many generations). Conversely, we view an intervention as “supplemental” when a given intervention has costs on a per-individual basis, and the benefits of that intervention do not persist over time. For example, outplanting of fragments will have a cost per fragment, and will boost population size, but will not change underlying vital rates of colonies in the population. Similarly, direct provision of superfoods may increase the survival, growth, and/or fecundity of those colonies that receive the intervention, but this intervention would need to be repeated in successive generations for benefits to be sustained. In between the above extremes, one can imagine interventions whose costs are highest at the initial implementation phase but have a lower but non-zero ongoing cost. Many traditional management approaches would fall into this category, where a change to a regulatory regime (for example, elimination of fishing methods that destroy habitat or remove herbivores) would carry a high initial cost of compensating individuals and businesses that utilized those methods, but on an ongoing basis would include (presumably lower) costs of ensuring compliance and potentially foregone economic benefits.

Theoretically, structural and supplemental interventions can produce similar outcomes, e.g. adding new individuals to the population each year can have the same demographic consequences as altering survival probabilities, because new individuals contribute to successful growth and reproduction. However, this supplemental strategy can be costly (Bayraktarov et al. 2019; Hughes et al. 2023), as those numbers must come from outside the population (e.g., a nursery or corals of opportunity or previously harvested fragments) and be actively deployed every year. A review of coral restoration in Indonesia found that 84% of projects reported a supplemental approach (i.e., some form of coral transplantation) and 16% relied on natural larval recruitment upon artificial reefs (Watt-Pringle et al. 2024). The two demographic approaches are not mutually exclusive; for example, genetically different or selectively bred corals can be supplementally added to potentially increase long-term (i.e., structural) survival and reproduction (Drury et al. 2022, Bay et al. 2017). Ultimately, demographic approaches are needed to assess the relative utility of supplemental versus structural modifications to population and their effectiveness for reef conservation.

In this paper, we will model “supplemental” approaches as one-off augmentations to the number of individuals in one or more life stages. This approach is intuitive for approaches such as deploying fragments onto the reef. However, we also adopt this approach for supplemental approaches that temporarily boost demographic rates, such as provision of superfoods. Our reasoning is that such interventions typically will have a cost per colony treated, and thus for a given investment, a fixed number of individuals will have higher growth, survival, and/or fecundity as the population grows or shrinks. Therefore, the net effect in the next year would be an increase of a fixed number of individuals in one or more life stages, rather than a change to the projection matrix that records per-capita contributions to the next generation or time step for the population as a whole.

Optimal restoration approaches will likely differ among geographic locations and coral taxa with different sensitivities to demographic change across their life cycle. Coral species exhibit an array of life history strategies because of energetic trade-offs among growth, survival and reproductive rates (Edmunds et al. 2014, McWilliam et al. 2023; Stearns 1989). Indeed, this array of strategies supports coexistence among species (Chesson 2000) and is essential for reef ecosystem function (Bellwood et al. 2019). For corals, major morphological groups (or growth forms) tend to reflect life history strategies (Jackson 1979; McWilliam et al. 2023; Dornelas et al. 2017). For instance, branching and tabular coral growth forms are specialized to compete for reef space via early and rapid growth (Baird & Hughes 2000), but typically have lower survival rates across life stages (Madin et al. 2016). Reproductive output varies dramatically among coral taxa, yet tends to be similar within growth forms, and highest in taxa which reach large colony sizes (Hall and Hughes 1996; Álvarez-Noriega 2016). Identifying life stages in which lambda is sensitive to change in specific vital rates is therefore necessary to design suitable conservation and restoration plans, acknowledging that these may vary across species.

In this study, we use demographic modeling to explore how changes to population structures can shift a population from decline to recovery for the purposes of coral conservation. We collated existing coral matrix population models capturing a range of geographic locations and coral morphologies (Table 2) to address a question that all reef resource managers and restoration practitioners should ask: what changes in transition probabilities or outplant numbers are needed to increase a population’s yearly growth by a given amount? We explicitly compare two approaches to reef conservation, the supplemental approach in which colonies are outplanted, and the structural approach in which vital rates are modified in a sustained fashion, quantifying the relative effort needed for each to affect population growth rates. We also examine the different life stages of coral in which changes in vital rates or abundances have the largest effects on population growth across different taxa. Our study highlights that demographic approaches are needed to inform decisions regarding the allocation of effort when attempting to preserve and restore coral populations.

## Methods

We collated 28 projection transition matrices, representing recruit, juvenile, and adult life history stages, spanning 22 species from six growth forms of reef-building corals (Table 2). Of these matrices, 11 were already three-by-three stage matrices representing recruits, juveniles and adults from the Comadre database (Salguero-Gómez et al. 2016). Only complete matrices that captured unmanipulated population growth were included from Comadre. Another 11 species matrices were generated from the Trimodal dataset (Lizard Island, Great Barrier Reef; Madin et al. 2023) by transposing yearly colony growth and survival data into three-by-three matrices using the stage size classifications in Table 2 (see Supporting Methods, Fig. S1 and Fig. S2). Six more species matrices were generated from NOAA’s Coral Reef Conservation Program-funded Vital Rate Project across the Hawaiian Archipelago (Rodriguez et al 2021). For those matrices that began as larger than three-by-three, we selected stage transition sizes using best-available, species-specific data to highlight stage-relevant transitions in mortality and reproduction. To set the recruit to juvenile stage size boundary, we employed the field-observed, size-dependent decline in mortality rates from the high mortality “recruit” stage to moderate “juvenile” stage (Fig. S1). To set the juvenile to adult stage boundary, we used best-available estimated size of first reproduction (Fig. S2). For 11 of the 17 models available as larger than three-by-three, this reproductive data was co-collected with demography. For the other 6, we used best available estimates for the Coral Trait Database (Madin et al. 2016). We assumed that the vital rates reflected in population matrices capture the average closed population over its entire geographic range.

To compare the demographic effects of restoration across coral taxa in wide array of demographic states, we assumed that all populations were at stasis when matrix data were collected—neither growing nor declining. To do this, we modified the recruitment rates of taxa (which were not recorded in field studies) and used a least squares function and the *optimize* function in R to estimate the amount of recruitment generated by adults that would result in a population with λ=1 (R Core Team 2025; Fig. S3A). Estimated levels of recruitment per adult colony required to make λ=1 were consistent with other levels reported in the literature (Madin et al. 2012; McWilliam et al. 2023). To estimate the sensitivity of results to the stasis assumption, we also estimated recruitment rates for populations assuming λ=0.9, 0.95, 1.05 and 1.1, which captures starting populations in decline through to growth (Supplemental Methods). These demographic models assume a closed stock-recruitment relationship, which, given the broad larval dispersal common among corals, requires that we assume that the resulting matrix represents vital rates covering the spatial scale of the demographically coupled population, likely at 10s of km.

We used the least squares function to estimate transition cell changes (structural) and outplant numbers (supplemental) required to achieve population growth increases of 1, 2, 5 and 10%. For the multiplicative approach, we increased a given transition cell value, while holding all other cells constant, until attaining levels of λ=1.01, 1.02, 1.05 or 1.1 (e.g., Fig. S3A). Cell values were allowed to increase above 1, which is theoretically the maximum transition probability for all cells that do not reproduce. However, we only report changes for the species for which cell values reflecting transition probabilities were one or less. We did not include the calculated changes for “shrinkage” cells, as the perturbation resulted in impossibly large values. That is, increasing population growth rate by elevating transitions from a large stage to a smaller stage can only occur when total survival probability in the resulting stage is above one, which can only be accomplished by adding individuals to the population, such as by reproduction.

For the supplemental approach, we increased each stable size structure (SS) value, while holding all others constant, until the summed matrix multiplication of the baseline transition matrix and the SS reached 1.01, 1.02, 1.05 or 1.1 (e.g., Fig. S3B). The amount of SS increase for a stage was multiplied by 1000 to express the number of outplants required per 1,000 individuals in the population. We also present these numbers in terms of the outplanted coral area per year (i.e., the planar area of coral required from an outside source), which was done by multiplying the number of individuals in each stage by the midpoint stage sizes in Table 2. We multiplied these values by 10,000 to express as outplant area per hectare (10,000 m^2^) of population cover. Note that supplemental metrics only reflect the levels required in the first year; that is, proportionally more additions are required as the population size increases assuming the absence of population density effects.

We estimated typical coral population sizes and population domain areas (i.e., the total area of living coral in a population) using NOAA’s National Coral Reef Monitoring Program (NCRMP) data on coral colony density across the US-affiliated central Pacific. The NCRMP effort includes a stratified random survey of reef sites across 37 islands which enumerates coral colonies, returning colony-level taxonomic identification, colony density, and survey sector reef area (Towle et al. 2022). From this data set, we highlight 80 taxa that represent the most-robust species-level identifications present in the dataset and then relied on the survey design estimates of sector-level colony number; that is, the estimated census population size at the sector scale (Smith et al 2011; Winston et al 2019). The NCRMP survey design assesses adult colony density in three depth-dependent strata per sector (0-6 m, 6-18 m, 18-30 m), so sector-level abundance (population size) for each representative species was calculated as the product of strata-level mean colony density and stratum area, summed across the three strata in each sector (Smith et al 2011; Winston et al 2019). Here, the sector represents roughly 10s of km of coastline (i.e., island scale for small islands [e.g., Jarvis, Pagan]) or side-of-island scale for larger islands (e.g., Oʻahu, Guam). Previous work on the modeled distribution of juvenile colony density, show an apparent stock-recruitment at sector scale, which suggests this scale is a reasonable proxy for a demographically coupled population (Couch et al 2023). While we are presenting projection matrices from three regions (Caribbean, Australia, Central US-Pacific), we only present the estimates of population sizes from the US Pacific.

## Results

Across the 28 population matrices, yearly survival probabilities tended to increase as colonies got larger (Fig. 2). Growth probabilities declined slightly with increasing size, and growth from recruit to adult (skipping the juvenile phase) had the lowest probability, and it was only observed in some massive and laminar species populations. Shrinkage (partial mortality or fission) to smaller stages tended to occur more for the adult stage. By way of example, Fig. 1A shows the matrix for *Acropora cytherea* and the blue transition cell represents the estimated number of successful recruits generated by an average adult colony per year; in this case, for the population to remain in stasis, only 8-9 of the hundreds of thousands of larvae generated by an adult colony must recruit back into the population recruit stage class (i.e., approximately 250 cm^2^ in area). The green cells show probabilities of surviving and growing to larger stages; yellow cells show probabilities of surviving and remaining at the same stage; and red cells show surviving and shrinking via partial mortality or fission to smaller stages. The vector to the right of the transition matrix in Fig. 1A is the population’s stable size structure (i.e., the transition matrix’s eigenvector): the proportion of colonies at each stage as defined in Table 2 once the population has stabilized (i.e., 76% of the colonies would be recruits, 17% juveniles and 7% adults; Caswell 2018).

**Figure 2.**
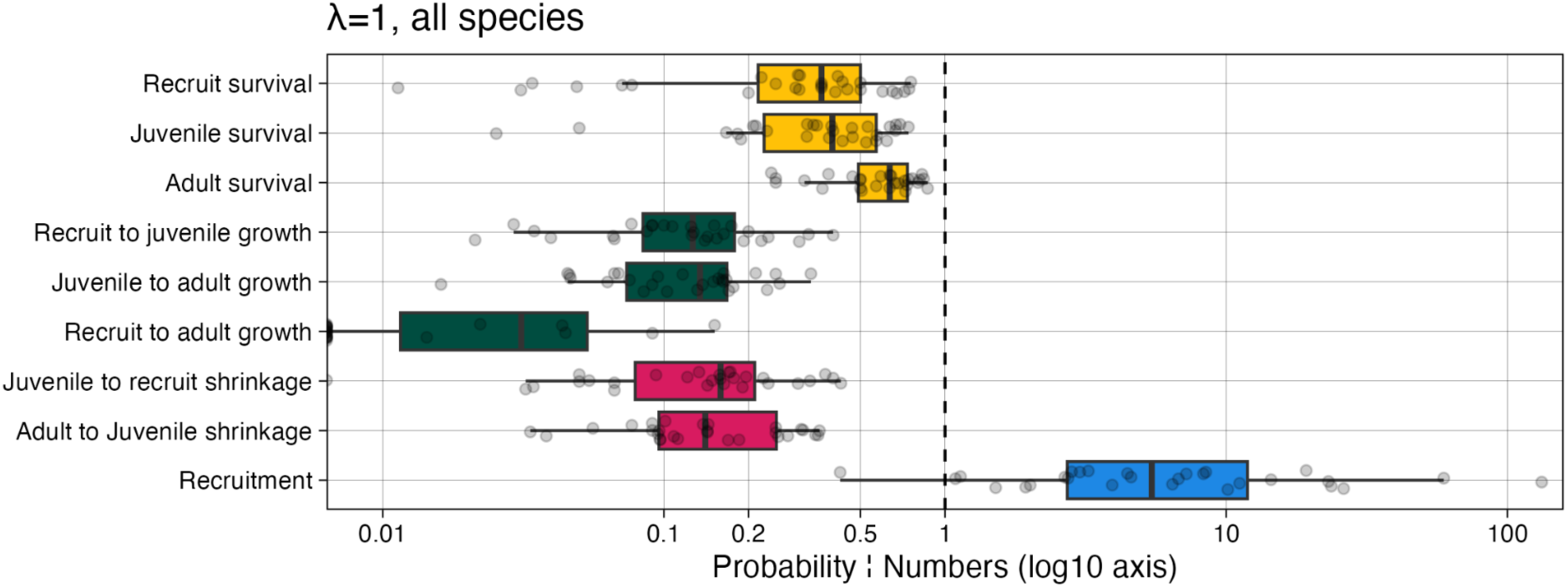
Boxplots for transition cells values for the 34 species matrices. The color scheme reflects that of Fig. 1 with survival probabilities in yellow, growth and survival probabilities in green, shrinkage in red, and contribution of adults to recruitment in blue.

Demographic models captured the changes in colony numbers and transition values needed to shift populations from stasis to growth. Using *Acropora cytherea* as an example, we find that increasing population size by 5% (equivalent to increasing λ to 1.05) would require supplementing the population with either 140 recruits, 64 juvenile or 5 adult colonies per 1000 colonies in a population per year (Fig. 1B). Similarly, for a summed planar area of 100 m^2^, we find that 3% of this area (3 m^2^) would need to be outplanted as new adult corals each year to increase population growth rate by 5%. Taking a structural approach, a 5% increase in λ can be achieved by increasing recruit survivorship from 0.25 to 0.6 (assuming all other transition probabilities remain the same; Fig. 1B), equivalent to a 2.4-fold increase in recruit survival rates (parentheses, Figure 1B). On the other end of the life-stage spectrum, increasing survival of adult colonies from 57% to 78% achieves a 5% increase in population growth, equivalent to a 1.36-fold increase in adult survival. Finally, 5% increases in population growth of *A. cytherea* can be achieved by increasing the survival of juvenile colonies that grow into adults and the probability of recruitment; each of these requiring an approximately 1.7-fold change.

Changes in transition values needed to achieve 5% increase in population growth varied across the 28 matrices in the analysis, though some consistencies emerged across different life histories and biogeographic regions (Fig. 3). Species for which required transition values became greater than one (Fig. 3A, red points) indicated that the multiplicative approach was not a viable option. For instance, improving adult survivorship in order to increase population growth rate by 5% was only viable for 79% of our populations. Considering only the population for which a strategy is possible, two transition cells—adult survival and recruitment—required the least change in value to achieve a 5% increase in population growth (Fig. 3B), which were also reflected in matrix elasticities for adult survival and recruitment that quantify the relative sensitivity of population growth to the various transitions (Fig. S4; Caswell 2018). Increases in successful recruitment or adult survival present the most consistent, demographically efficient means to increase population growth, with the caveat that the adult survival approach is only viable for 23 of the 28 populations. The small changes in recruitment and adult survival needed to boost population growth were largely consistent across coral growth forms (Fig. 3C). Changes required for other transition cells were higher and more variable, reflecting the sensitivity of growth forms to different changes in demographic rates. For example, the low growth in massive taxa and the low juvenile survival in laminar taxa required higher necessary changes in transition probability to boost population growth (Fig 3c; Fig S5). Meanwhile, higher recruit survival in massive taxa meant that smaller shifts in recruit survival were needed relative to other growth forms (Fig 3; Fig S5). The small changes in recruitment and adult survival needed to boost population growth were also consistent across biogeographic regions (Fig 3D) with the exception that the Caribbean showed greater efficacy (lower transition multiplier) of elevated recruitment survival for some species, but not all as reflected by high levels of variation.

**Figure 3.**
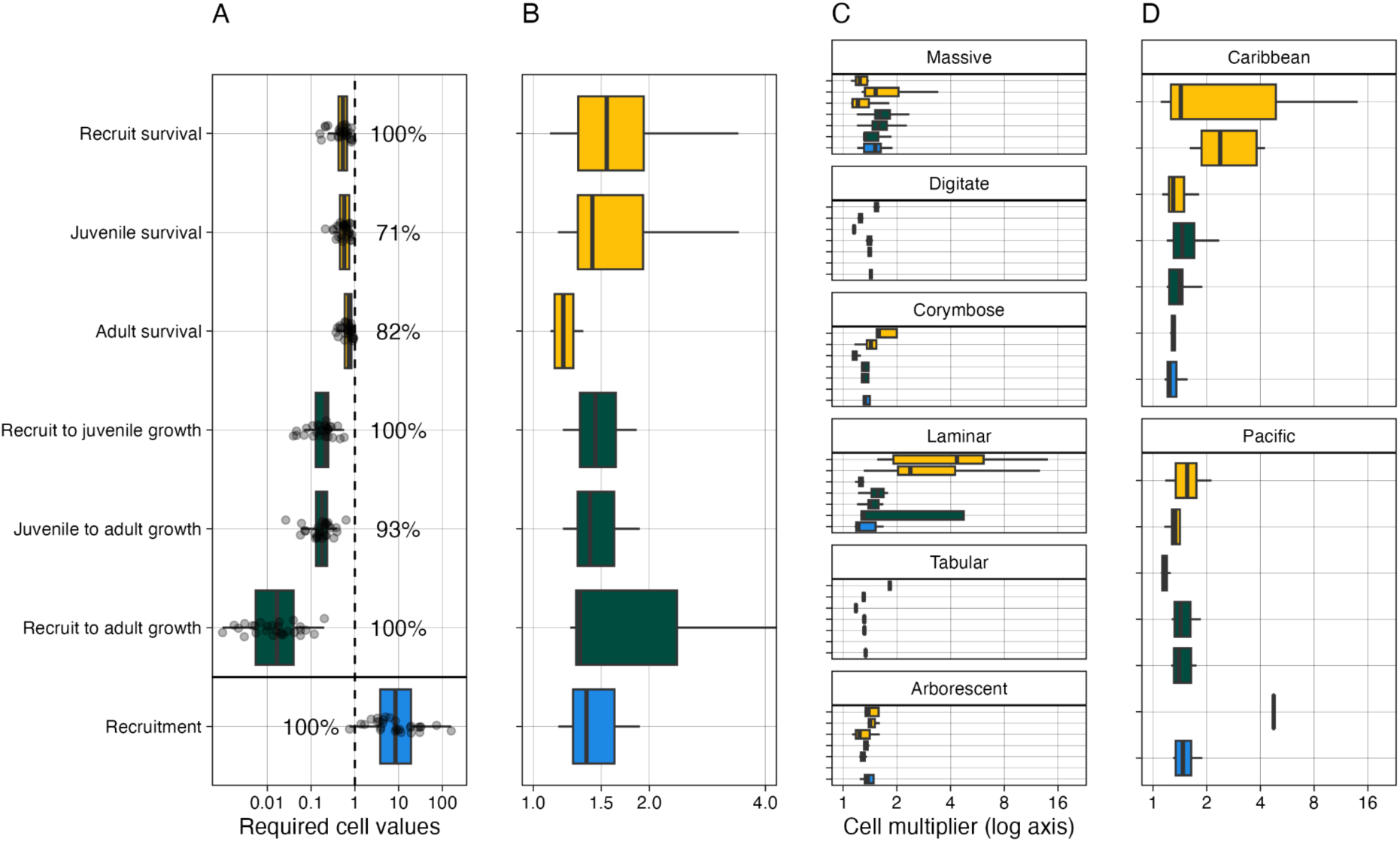
The structural approach. (**A**) The cell values required to increase population growth rate by 5% (λ=1.05) summarized for all species. Transition values cannot exceed one for cells that do not generate individuals unless individuals are artificially added. The percent of species for which changing cell values is possible are given. (**B**) The proportional change in cell values required to increase population growth rate by 5%, only for species for which the approach is possible. Proportional change is also shown for (**C**) species growth form and (**D**) biogeographic region. The color scheme reflects that of Fig. 1 with survival probabilities in yellow, growth and survival probabilities in green, and contribution of adults to recruitment in blue.

Increasing population growth by supplementing corals required approximately 100 recruits or juveniles per 1000 colonies in the population in the first year (Fig. 4A, left), or 5–10 adults; and this number would increase proportionally as the population size increased. These numerical additions translate into similar amounts of live coral area for recruits and adults of approximately 2–5 m^2^ yearly for every 100 m^2^ of adult coral area in a population (Fig. 4A, right). In contrast, juveniles require approximately an order of magnitude more coral material since they require large numbers of corals (comparable with recruits) but with substantially increased areas (Table 2). These results were consistent among growth forms, with notable deviations for massive and laminar growth forms. For example, significantly more recruits or juveniles would be required relative to adults for the laminar populations due to low early survival rates, suggesting that a strategic focus on adults may more efficiently elevate population growth. Conversely, fewer recruits would be required to elevate population growth for massive species relative to other growth forms, likely because of higher rates of recruit survival (Fig. 2). Desired levels of population growth rate scale approximately linearly with the amount of coral material added to the population each year (Fig. 4B). For instance, for a population size of 1000 individuals, adding on the order of one adult per year is estimated to increase population growth by 1% and adding on the order of 10 adults would increase it by 10%.

**Figure 4.**
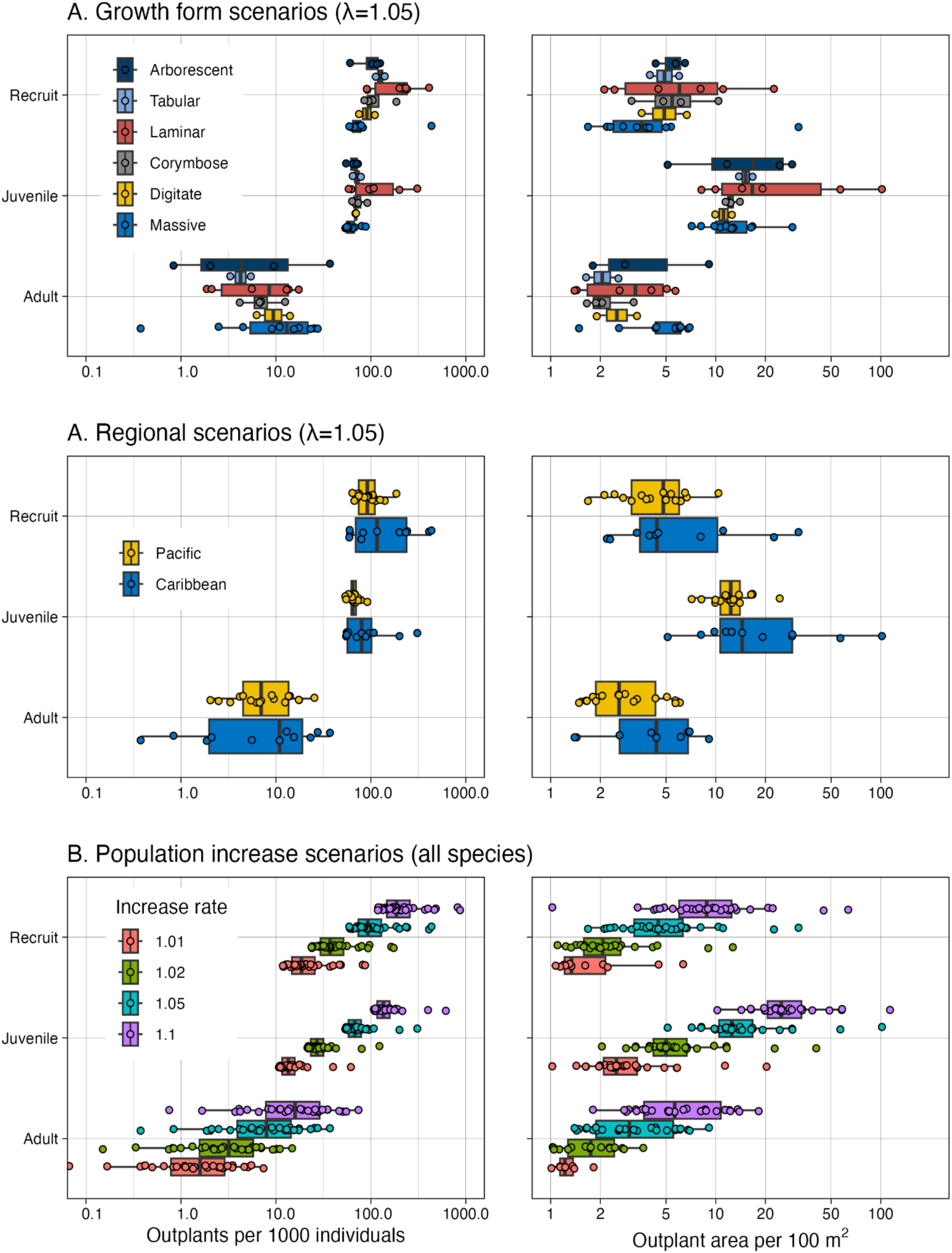
The numbers approach. (**A**) Boxplots of the number of outplants (left) and outplant area (right) for each life history stage required to increase yearly population growth rate by 5% (λ=1.05), summarised by growth form and region. (**B**) Boxplots of the number of outplants per 1000 individuals (left) and outplant area per 100 m^2^ of population area (right) for each life history stage required to increase yearly population growth rate by 1, 2, 5, and 10% for all species.

Both supplemental and structural restoration approaches depend on the scale of implementation (Fig. 5). The number of yearly adults that need to be outplanted to achieve a 5% increase in population growth differs by two orders of magnitude across species (Fig. 5B). Assuming that sector-scale population sizes seen across the US-affiliated Pacific are reasonable estimates for other populations and based on mean estimates of census population size (∼625,000 individuals, Fig. 5A), a minimum of 160 (*Montastrea annularis*) and a maximum of 26,000 (*Acropora cervicornis*) adult colony outplants per year are necessary to alter the dynamics of a population. Orders of magnitude greater numbers of additional recruits are necessary to alter population growth by 5%, with a minimum of 33,000 and a maximum of 630,000 recruits added to the natural rates every year, based on the average population size in the dataset (Fig. 5C). Such numbers highlight the difficulty of altering population growth at large (regional) scales using the supplemental approach.

**Figure 5.**
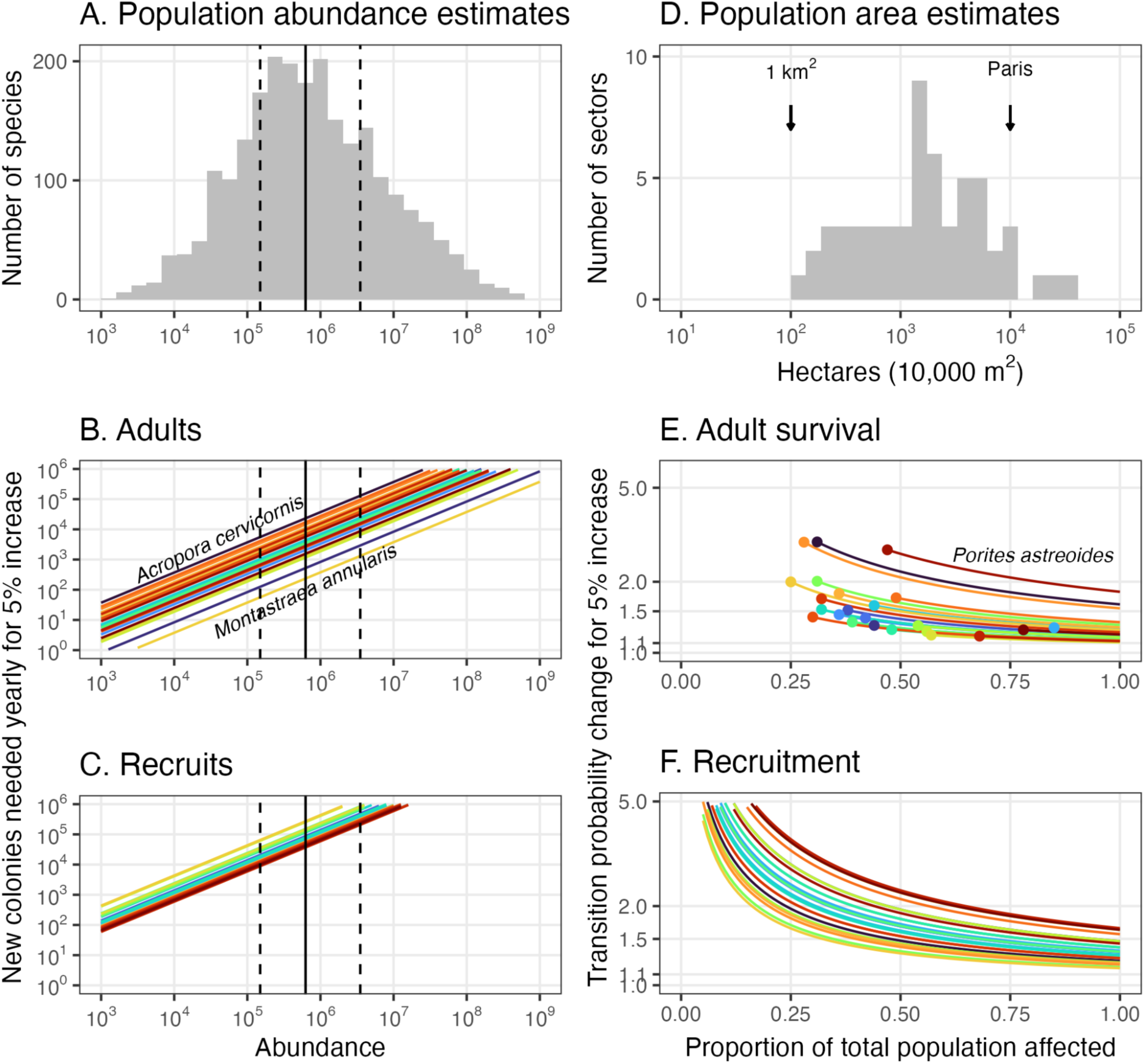
(**A**) Population abundance for 80 species estimated from NOAA NCRMP Pacific benthic survey data. Comparison of approaches relative to species population sizes. (**B**) The number of yearly adult outplants required to increase population growth rate by 5%, with each colored line representing one of the 34 examined transition matrices. The species at either end of the range are labeled. (**C**) The number of yearly recruit-sized outplants needed to increase population growth by 5% for each of the 34 matrices. (**D**) Population-scale reef area estimates from NOAA Pacific benthic survey data. (**E**) The change in adult survival probability needed to increase population growth rate by 5% when considering the proportion of the population affected by a conservation measure for each of the 34 matrices. Points indicate the theoretical limit; that is, where adult survival probability is 100%. (**F**) The change in recruitment probability needed to increase population growth rate by 5% for each of the 28 matrices.

Elevating adult survival probability is also likely to be most effective if the benefit can be implemented across the entire population (Fig. 5D). However, when we model a realistic intervention that does not benefit the entire coral population, we can show that the vital rate increases required to meet a 5% shift in population growth rate (lambda) are relatively stable when only subsets of the population are affected (Fig. 5D-E). Indeed, for many taxa the necessary improvement to survival remains stable (< 1.5-fold increase) as long as > 50% of the population is affected (Fig. 5E). Similarly, a five-fold increase in recruitment was sufficient to shift lambda to 1.05 for all taxa as long as > 25% of the population was affected by the changes (Fig. 5F). Nevertheless, the number of additional recruits required becomes exceedingly high if small portions of the population are impacted (Fig 5F). Moreover, we identify a critical threshold in survival beyond which a 5% change in lambda is theoretically impossible because survival has already reached 100% (points in Fig. 5E). For many taxa, this threshold occurs between 25-50% of the population affected, demonstrating the importance of capturing large proportions of the population to achieve meaningful shifts in lambda, a feat that requires conservation action to be implemented across large areas (Fig 5D).

## Discussion

Our analysis suggests that modifying vital rates at critical stages of the life cycle, using interventions whose impacts are sustained, is the most effective way to grow coral populations. This “structural” approach focuses on modifying factors that influence long-term survival, growth and recruitment, and shows considerable promise if it can be applied in ways that benefit large portions of the effective population (Fig. 5D-F). A wide range of conservation measures can be employed to modify vital rates (Table 1), yet demographic models suggest that those which affect adult survival and recruitment rates are likely to be the most successful because they require the least change. If applied to the whole population, a 1.2-fold increase in survival (20% increase) or a 1.5-fold increase in recruitment (50%, on average) was sufficient to boost population growth by 5%. In contrast, relying solely on outplanting additional colonies to modify coral population growth rate—the supplemental approach—is likely to be inefficient for most species, particularly those with high abundances (Fig. 5A-C). While it was possible to boost population growth by adding colonies, we find that yearly additions of 5-10% of the existing population size in recruit form, or 0.5-1% of the existing population size in adult form, were necessary to alter population trends (Fig. 4). Genetic models suggest that a similar level of adult transplantation at similar orders of magnitude (2-5% of the population per year) could prevent extinction, but not population decline, under moderate emissions scenarios (formerly RCP6.0) only if transplanted corals were pre-adapted to warming conditions (Bay et al. 2017). Depending on the population size, the number of new colonies needed can be exceedingly large (Fig. 5A-C).

**Table 1.** Common approaches for coral restoration and conservation and their demographic approach.

| Activity | Demographic Approach | Description | Examples |
| --- | --- | --- | --- |
| Allee Effect Mitigation | Supplemental<br>Structural if successful | Increasing density beyond the minimum required for successful fertilization. | Ricardo et al. 2025 |
| Artificial habitat | Structural | Deployment of artificial structures for attracting coral and other biodiversity with a focus on the recruitment stage. | Knoester et al. 2023; Reichert et al. 2025; Tanaya et al. 2025 |
| Assisted evolution | Structural | A range of interventions that speed up the natural process of coral adaptation that might improve growth and survival. | Van Oppen et al. 2015; Caruso et al. 2021; Drury et al. 2022; Baker et al. 2025 |
| Assisted migration | Supplemental<br>Structural if successful | The delivery of corals with environmental tolerances to another based on anticipated environmental changes. | Van Oppen et al. 2014; Caruso et al. 2021; Baker et al. 2025 |
| BioRock Electric Reef | Supplemental | Like substrate stabilization, but with the addition of low voltage electricity applied to metal mesh to promote calcification. | Goreau & Prong 2017; Goreau 2022 |
| Coral gardening | Supplemental | The asexual propagation of corals onto a reef typically following a nursery phase. | Page et al. 2018; Tortolero-Langrancia et al. 2020 |
| Coral repositioning | Supplemental | Securing dislodged corals back onto the reef. In some cases, corals will be fragmented before being repositioned. Like coral gardening and transplantation, but without a nursery phase. | Garrison & Ward 2008; Nava & Figueroa-Camacho 2017; Williams et al. 2019; Ladd et al. 2019; Sam et al. 2021 |
| Coral seeding | Supplemental | The settlement of coral larvae or attachment of small fragments onto structures for reef deployment. | Forsman et al. 2015; Lindahl 2003; Suzuki et al. 2011; Williams et al. 2019; Ladd et al. 2019 |
| Coral superfoods / probiotics | Supplemental | Administering nutrients or supplementing coral microbiomes to boost survival and growth. | Doering et al. 2021; Peixoto et al. 2017 |
| Coral Transplantation | Supplemental<br>Subtractive from source population | Like assisted migration or coral seeding, corals are moved from one place to another, but environmental tolerances are not considered, and adult colonies are generally used. Supplementing existing populations with individuals destined for mortality due to development, without a nursery phase. An umbrella term that can also include coral gardening and repositioning. | Lindahl 2003; Forrester et al. 2019; Kikuzawa et al. 2021 |
| Early maturity | Structural | Applying artificial selection (e.g., breeding, selectively targeted out-planting) to promote maturity at smaller sizes. | Randall et al. 2020 |
| Herbivore fishing restrictions | Structural | Decreasing competition for space on the reef and improving recruitment, growth and survival of coral. However, recruitment can be hindered by herbivores in the absence of habitat complexity. | Penin et al. 2011; Bonaldo et al. 2017<br>Morais & Bellwood 2018; Strain et al. 2019 |
| Larval-based restoration | Supplemental | Supplying large numbers of coral larvae to damaged reef areas. | Heyward et al. 2002; Edwards et al. 2015; Dela Cruz & Harrison 2017; Doropoulos et al. 2019; Harrison et al. 2021 |
| Macroalgae removal | Supplemental if by people<br>Structural if sustained by herbivory | Suppress macroalgae proliferation to facilitate coral recovery. In some cases, paired with urchin releases. | Neilson et al. 2018; Smith et al. 2022 |
| Pollution & sedimentation mitigation | Structural | Generally improving the environment improves all demographic rates. | Donovan et al. 2021; Donovan et al. 2020; Gove et al. 2023 |
| Spawn capture and rearing | Supplemental | Collecting reproductive gametes during spawning events for culturing. | Heyward et al. 2002; Doropoulos et al. 2019; Dela Cruz & Harrison 2020 |
| Substrate stabilization | Structural | Laying an artificial substrate (e.g., mesh) over areas of unconsolidated reef rubble. | Fox et al. 2019; Ceccarelli et al. 2020 |
| Urchin releases | Structural | Like herbivore fishing restrictions, but potentially more disruptive to coral recruitment. | Neilson et al. 2018; Barrows et al. 2023 |

The population size affected by a conservation initiative strongly influences both the structural and supplemental approaches, which ultimately depend on the spatial scale of conservation. At the likely scale of demographically connected populations (i.e. stock-recruitment scale) we estimate the census population sizes of most coral species to be in the range of 10^5^ to 10^7^ individuals (Fig. 5A). These island-scale values are lower than many Pacific coral population estimates, as estimates that span full species ranges suggest that roughly two-thirds of species have population sizes exceeding 100 million colonies and one-fifth exceeding a billion colonies (Dietzel et al. 2021). However, recent taxonomic revisions of reef corals based on molecular data are showing that our ability to delineate species is poor and that what are considered large populations are actually made up of multiple species with smaller population and geographic ranges (Bridge et al. 2024, Rassmussen et al. 2025). Quantifying population growth at this more local scale is challenging, and more precise estimates would require information on dispersal dynamics and spatial differences in vital rates (Hunter & Caswell et al. 2005). Nevertheless, our simplified extrapolation of population growth suggests that outplanting as few as 500 adult colonies per year for rare species, or as many as 50,000 for common species, was necessary for a 5% increase in population growth at the scale of the NCRMP sectors (Fig. 5B-C). This result demonstrates the challenges of this strategy for typical coral population sizes unless a substantial amount of coral material is available, or the population is small or limited to an area with high levels of self-recruitment, like an embayment. In contrast, structural reef conservation measures, which focus on enhancing vital rates, can often effectively capture large components of a coral population. For example, networks of Marine Protected Areas can cover a large geographic space (> 1000 hectares; Selig & Bruno 2010; Smith et al. 2025) and thus boost coral vital rates (e.g., improved herbivory and coral recruitment) across a large portion of a coral population. Similarly, reducing greenhouse gas emissions is likely to benefit most, if not all, of a coral population by reducing mortality from climate stressors, such as recurrent bleaching (Hughes et al. 2023), although the benefits of greenhouse gas reductions may take considerable time to be realized.

For local interventions, structural approaches also offer the greatest chance of boosting coral population growth at the site or reef scale, especially if they limit the need for costly yearly intervention. For example, artificial structures can enhance local recruitment by providing settlement substrate, modifying water flow, and increasing larval settlement probabilities (Nguyen et al. 2022; Higgins et al. 2022; Fig. 6), although longer-term studies have shown restoration success to be highly site dependent (Fox et al. 2019). Our demographic analyses begin with the smallest recruit size classes reported in published matrix models rather than newly settled spat, reflecting a broader limitation of the available demographic literature. Consequently, our calculations assume that increases in settlement ultimately translate into proportional increases in the abundance of recruits surviving to these modeled size classes. Under this assumption, if even a small fraction of incoming larvae experience a 10-fold or greater increase in settlement success (as has been shown possible; Fig. 6), the resulting increase in recruitment would be more than sufficient to increase local population growth by 5%, at least until density dependence begins to increase mortality. Interventions that improve both settlement and early post-settlement survival would be expected to strengthen this demographic effect. Coral populations on reefs can still be vast (e.g., a 1-hectare reef with one coral per m² has a population of 10,000), thus requiring hundreds of new recruits or multiple adult colonies to be added every year to modify population growth, even at a relatively small scale. Ultimately, boosting local population growth depends on recruitment, which is highly variable through time and space owing to the open nature of coral populations. Understanding this variability and source-sink metapopulation dynamics will be crucial for a demographically informed approach to coral conservation (Crowder et al. 2000; Hansen 2011).

**Figure 6.**
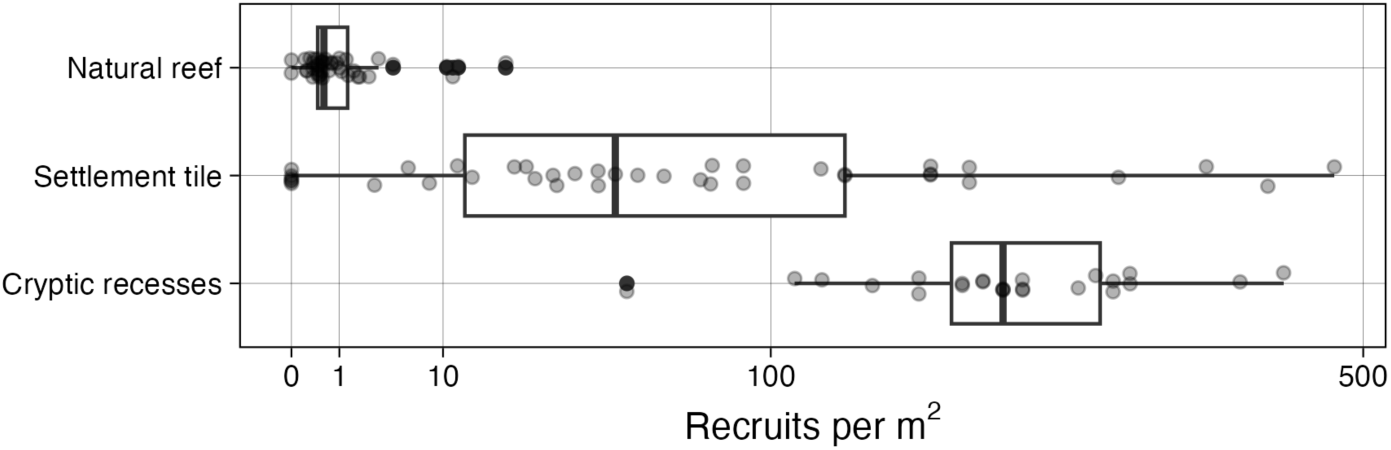
The density of coral recruits on natural reefs (Reichert et al. 2025), traditional settlement tiles (Bramanti and Edmunds 2016; Edmunds 2022; Brown 2004; Friedlander and Brown 2005), and structures designed with cryptic recesses (Doropoulos et al 2016; Reichert et al. 2025). See supplemental methods for details.

We find two critical demographic transitions whereby small changes had strong effects on population growth. First, adult survival universally required the least change in probability to increase population growth. This result was expected, given that corals produce many offspring that mostly perish (Type III survivorship; Crouse et al. 1987; Tsounis & Edmunds 2016). For example, a 20% increase in average adult survival can enhance population growth by 5% (Fig. 2B), which is remarkably consistent across growth forms and regions (Fig. 2C,D). Past studies suggest that focusing on adult survivorship provides the most efficient use of resources, delivering significant population benefits for smaller interventions. For instance, protecting mature female turtles from trawling led to population increases in the endangered loggerhead turtle (Crouse et al. 1987). However, our analysis suggests that this approach is not viable for all species populations, with 21% of the populations requiring increases in survival surpassing 100% (Fig. 3A; Supplementary Table 1). Furthermore, increasing adult survivorship across large areas for sessile corals is complex, and would typically involve addressing environmental stressors like thermal anomalies or mitigating local stressors (Ateweberhan et al. 2013; Good & Bahr 2021; Donovan et al. 2021; Gove et al. 2023). Increasing adult survivorship is also an important focus of many restoration strategies (Table 2), including substrate stabilization (Fox et al. 2019; Ceccarelli et al. 2020), macroalgal removal (Neilson et al. 2018; Smith et al. 2022), artificial evolution (Van Oppen et al. 2015; Caruso et al. 2021; Drury et al. 2022), and artificial migration (Van Oppen et al. 2014; Caruso et al. 2021).

**Table 2.** A summary of the 28 matrices by species. Each life history stage shows the range of planar areas for each stage. In parentheses are the rough midpoints of each stage’s planar colony area, which is used to estimate colony areas in the modeling. ^a^Mercado-Molina et al. (2015); ^b^Madin et al. 2023; ^c^Hughes & Tanner 2000; ^d^Rodriguez et al 2021; ^e^Hernández-Pacheco et al. 2011. Growth form was collected from the Coral Trait Database (Madin et al. 2016).

| Species | Region | Matrices | Recruit (cm<sup>2</sup>) | Juvenile (cm<sup>2</sup>) | Adult (cm<sup>2</sup>) | Growth form |
| --- | --- | --- | --- | --- | --- | --- |
| *Acropora cervicornis*<sup>a</sup> | Caribbean | 2 | 0-100 (50) | 100-1000 (550) | >1000 (1500) | Arborescent |
| *Acropora cf. digitifera*<sup>b</sup> | GBR | 1 | 0-150 (75) | 150-300 (225) | >300 (375) | Digitate |
| *Acropora cytherea*<sup>b</sup> | GBR | 1 | 0-500 (250) | 500-2000 (1250) | >2000 (3000) | Tabular |
| *Acropora humilis*<sup>b</sup> | GBR | 1 | 0-200 (100) | 200-300 (250) | >300 (350) | Digitate |
| *Acropora hyacinthus*<sup>b</sup> | GBR | 1 | 0-200 (100) | 200-1000 (600) | >1000 (1400) | Tabular |
| *Acropora intermedia*<sup>b</sup> | GBR | 1 | 0-150 (75) | 150-500 (325) | >500 (675) | Arborescent |
| *Acropora millepora*<sup>b</sup> | GBR | 1 | 0-150 (75) | 150-300 (225) | >300 (375) | Corymbose |
| *Acropora nasuta*<sup>b</sup> | GBR | 1 | 0-150 (75) | 150-300 (225) | >300 (375) | Corymbose |
| *Acropora robusta*<sup>b</sup> | GBR | 1 | 0-200 (100) | 200-500 (350) | >500 (650) | Arborescent |
| *Acropora spathulata*<sup>b</sup> | GBR | 1 | 0-150 (75) | 150-700 (425) | >700 (1050) | Corymbose |
| *Agaricia agaricites*<sup>c</sup> | Caribbean | 2 | 0-20 (10) | 20-100 (60) | >100 (150) | Laminar |
| *Goniastrea pectinata*<sup>b</sup> | GBR | 1 | 0-20 (10) | 20-200 (110) | >200 (300) | Massive |
| *Goniastrea retiformis*<sup>b</sup> | GBR | 1 | 0-20 (10) | 20-200 (110) | >200 (300) | Massive |
| *Helioseris cucullata*<sup>c</sup> | Caribbean | 2 | 0-20 (10) | 20-100 (60) | >100 (150) | Laminar |
| *Montipora capitata*<sup>d</sup> | Hawaii | 1 | 0-20 (10) | 20-100 (60) | >100 (150) | Laminar |
| *Montipora patula*<sup>d</sup> | Hawaii | 1 | 0-20 (10) | 20-100 (60) | >100 (150) | Laminar |
| *Orbicella annularis*<sup>b,e</sup> | Caribbean | 4 | 0-50 (25) | 50-150 (100) | >150 (225) | Massive |
| *Pocillopora meandrina*<sup>d</sup> | Hawaii | 1 | 0-150 (75) | 150-700 (425) | >700 (1050) | Corymbose |
| *Porites astreoides*<sup>a</sup> | Caribbean | 1 | 0-50 (25) | 50-150 (100) | >150 (225) | Massive |
| *Porites lichen*<sup>d</sup> | Hawaii | 1 | 0-50 (25) | 50-150 (100) | >150 (225) | Massive |
| *Porites lobata*<sup>d</sup> | Hawaii | 1 | 0-50 (25) | 50-150 (100) | >150 (225) | Massive |
| *Porites lutea*<sup>d</sup> | Hawaii | 1 | 0-50 (25) | 50-150 (100) | >150 (225) | Massive |

Recent work on assisted evolution demonstrates that coral thermal tolerance can be improved through strong, multi-generational selection, potentially increasing survival-related vital rates under future environmental conditions (Lachs et al. 2026). From a demographic perspective, successful assisted evolution represents a structural intervention because improved genotypes may spread through populations and generate sustained changes in survival and reproduction over multiple generations. Our synthesis suggests that relatively modest improvements in recruitment and survival (∼20%) could substantially enhance population growth if those benefits are realized across large fractions of a population. The key challenge therefore is not whether genetically based interventions can, in principle, alter population dynamics, but whether gains demonstrated under experimental conditions translate into consistent demographic benefits under natural reef conditions and at ecologically meaningful scales.

Increasing recruitment success was the second leverage point for enhancing population growth (Fig. 3B). Despite the impressive fecundity of corals, recruitment and post-recruitment processes represent a marked bottleneck for coral population growth (Arnold & Steneck 2011; Ritson-Williams et al. 2009; Tsounis & Edmunds 2016; Doropoulos et al. 2021). We find that population growth can be consistently increased by 5% with a less-than 2-fold improvement in recruitment. Recruitment operates differently to other life stage transitions, because it is a multiplier of population numbers, rather than a probability of transition for a single individual, and so multiplicative changes are not bounded by one and so they are potentially easier to manipulate via management actions (Randall et al. 2020, Harrison et al. 2021). It is important to note that recruit size classes in our analysis are considerably larger than the early benthic stages typically considered in settlement ecology and restoration practice (Table 2). Nonetheless, given the large potential for larvae supply, capturing just a small fraction of wasted larvae has extraordinary potential for scalable restoration under structural intervention approaches and the assumption that probabilistic bottlenecks operate similarly from the settlement to recruit size stages (i.e., if density dependent effects can be avoided). Spawn capture and nursery-rearing of recruits is promising (Dela Cruz & Harrison 2020), but still faces the same limitations as coral gardening, relying on consistent outplanting to maintain its efficacy (Table 1). However, improving the passive capture of even a small fraction of wasted larvae has extraordinary potential for restoration. For instance, by conservatively improving natural settlement by just 2-fold and assuming sufficient space to avoid density dependent mortality, population growth could be increased by 10%. Reef managers could achieve such improvements by enhancing environmental conditions, such as reducing sedimentation or contaminants, promoting herbivorous fishes and urchins, restoring suitable habitats, stabilizing unconsolidated hard substrate, or deploying settlement substrates for coral larvae (Table 1). For instance, data from settlement structures with cryptic recesses routinely demonstrate improvements of 50-200-fold over natural reefs (Doropoulos et al 2016, Reichert et al.2025; Fig. 6). While density dependent processes (i.e., crowding and competition) act to thin these early life gains, the chances that increased numbers of juveniles make it through the early life bottleneck increases, sometimes dramatically (e.g., 20-50-fold after one year; Reichert et al 2025).

This paper considered various approaches to coral restoration from demographic first principles. By collating simple population matrix models and evaluating the influence of structural and supplemental demographic interventions, our analysis highlights the salient features of current restoration approaches and their likely effectiveness. As with all meta-analyses, several important caveats should be considered when interpreting these results, including differences in methodology, parameterization, and data quality among studies. For example, many older demographic matrices were already simplified into three-by-three stage structures and may not accurately represent recruit, juvenile, and adult size classes (Hughes & Tanner 2000 Hernández-Pacheco et al. 2011, Mercado-Molina et al. 2015, Salguero-Gómez et al. 2016). To facilitate comparison across studies, continuous demographic datasets were therefore converted into equivalent three-stage matrices based on survival and reproductive characteristics intended to approximate the structure of the older datasets (Rodriguez et al 2021, Madin et al. 2023). Our methods highlight the inconsistent definition of the recruit stage, which is larger in this study than other definitions and should be treated cautiously as mentioned above. Additionally, the simplified population projection matrix framework used here does not capture potentially important nonlinear demographic processes, such as density dependence, fertilization limitation, or Allee effects, and therefore the magnitude of supplemental versus structural intervention outcomes should be interpreted cautiously. Our analysis also assumed that transition matrices reflected populations approximately at demographic stasis. For declining populations, the intervention effort required to restore persistence is likely greater than the 5% increase in population growth modelled here. Nonetheless, our results appear remarkably insensitive to initial population growth rate (Fig. S6), suggesting that the relative advantages of different restoration approaches are broadly similar across both declining and growing populations.

There are contexts in which supplemental restoration at achievable scales can improve population outcomes. One subset of these would be locations where small, restricted populations are of high value, which minimizes the absolute scale of the supplemental intervention required and may justify an ongoing cost. Another critical example is where a supplemental action can have a structural effect, like the case of a population suffering Allee effects. In these cases, the supplemental intervention may push a population across a threshold above which demographic rates structurally shift into a new, more favorable regime. However, in many coral populations, raising population growth rates above replacement by outplanting more colonies into declining populations will require efforts at scales unlikely to be cost-effective. On the other hand, our demographic analysis suggests that structural efforts to the rates of survivorship and recruitment through combinations of traditional management and process-focused active interventions are likely the more effective strategy for responding to declining coral populations. Every coral conservation effort is unique and context-dependent, but we hope our demographic framework will provide resource managers and restoration practitioners a framework to understand and predict how restoration efforts will translate into meaningful outcomes.

## Authors’ contributions

Joshua Madin conceived the ideas and designed methodology; Joshua Madin, Mollie Asbury, Guanyan Keelung Chen, Jon Ehrenberg, Hendrikje Jorissen, Allison Nims, Jessica Reichert, Lomani Rova, Nina Schiettekatte and Devynn Wulstein participated in the workshop that conceptually developed the ideas; Joshua Madin, Thomas Oliver, Mike McWilliam and Hendrikje Jorissen collated the data; Joshua Madin and Thomas Oliver analyzed the data; Joshua Madin, Thomas Oliver and Mike McWilliam led the writing of the manuscript. All authors contributed critically to the drafts and gave final approval for publication.

## Statement on inclusion

Our study was a global review and was based on a meta-analysis of secondary data rather than primary data. As such, there was no local data collection. However, the geographical distribution of the authorship team broadly represents the major regions of interest in the meta-analysis and ensuring the appropriate interpretation of data and results from each region.

## Acknowledgements

The concept for this paper stemmed from a Geometric Ecology Lab workshop on disruptive science supported by the Hawaiʻi Institute of Marine Biology Director Council’s Innovation Fund.

## Data and Code

Data and code are available at https://github.com/jmadinlab/demographic-insights

