## Supplemental Material for "Demographic insights for coral restoration"

**Supporting Methods**. A key uncertainty with the matrices used in this study were the transition boundaries between recruits, juveniles and adults. Boundaries for the Comadre matrices were set during the design of the original studies and these cannot be changed. Survival rates in the recruit stage were consistently lower than the juvenile stage (Fig. 2), and the boundary between juvenile and adult was in the vicinity of reported mature colony sizes in the literature (Table 1). For the Trimodal species we can fine-tune the boundaries based on fecundity and survival data. For instance, we defined the recruit-juvenile boundary as the colony size where survival rate stabilized (Fig. S1) and the juvenile-adult boundary as the colony size where approximately 75% of colonies were reproductive (Fig. S3).

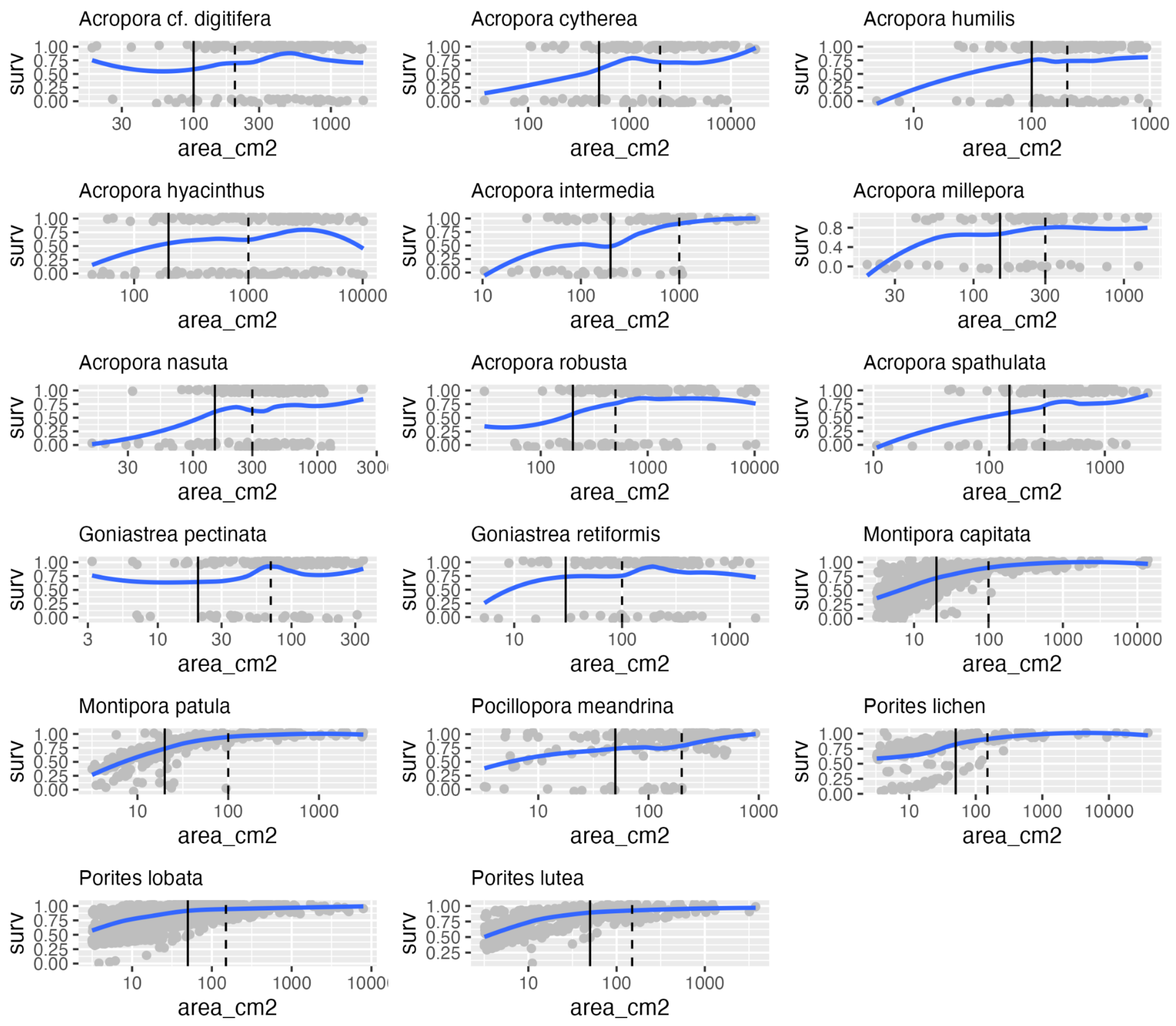


**Figure S1**. Yearly survival probability for Trimodal (Madin et al. 2023) and NOAA (Rodriguez et al 2021) species as a function of colony size showing transition boundaries used for recruit-juvenile (solid vertical lines) and juvenile adult (dashed lines corresponding with Fig. S2). Blue lines are loess smoothers to illustrate underlying size-survival patterns.

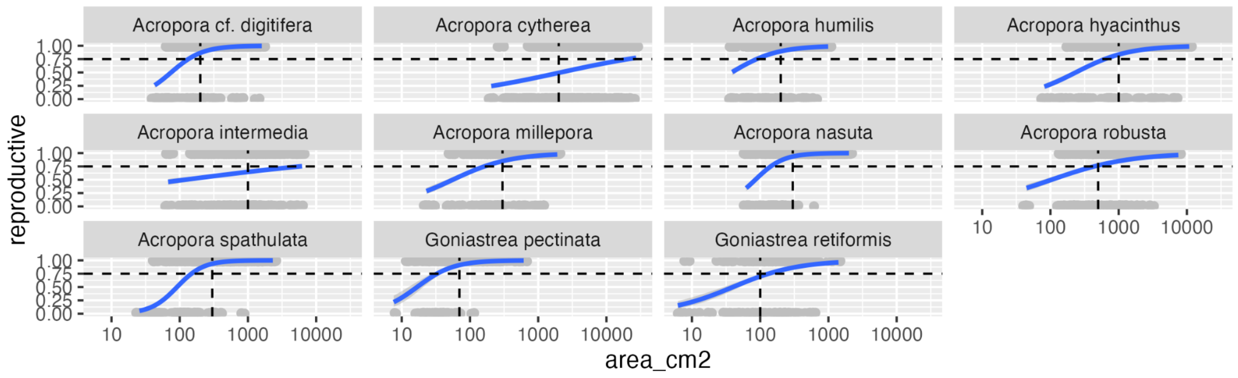


**Figure S2**. Size at reproduction for the 11 trimodal species (Madin et al. 2023) with logistic model fits (blue lines). We used the approximate size at which 75% of colonies are reproductive as the juvenile-adult boundary for transition matrices.

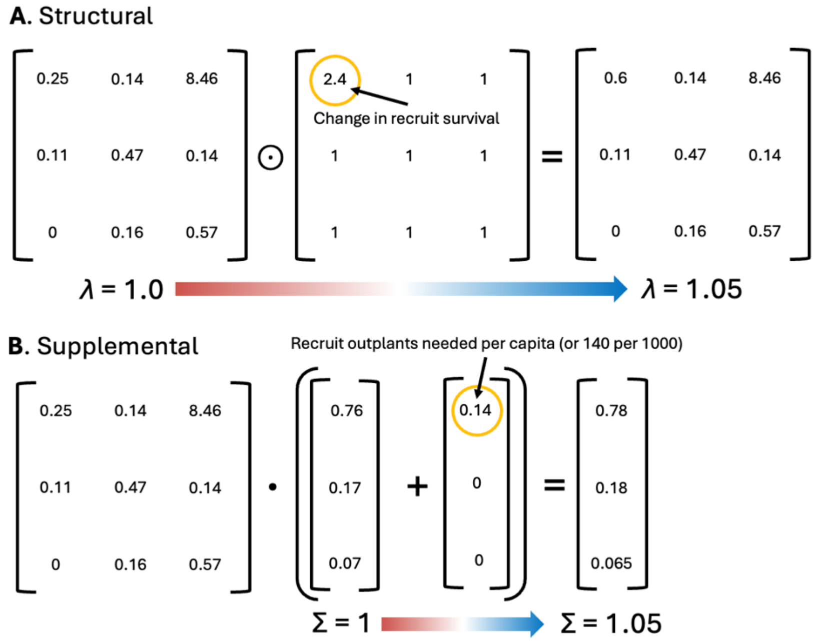


**Figure S3**. A demonstration of the matrix operations for (**A**) structural and (**B**) supplemental approaches to increasing year on year population growth by 5% for *Acropora cytherea*. ⊙ represents the Hadamard product and • represent the matrix product. Σ represents the sum. This example represents best estimates of recruit survival change in A and yearly recruit outplanting in B. The values circled in yellow were found using least square optimization in R.

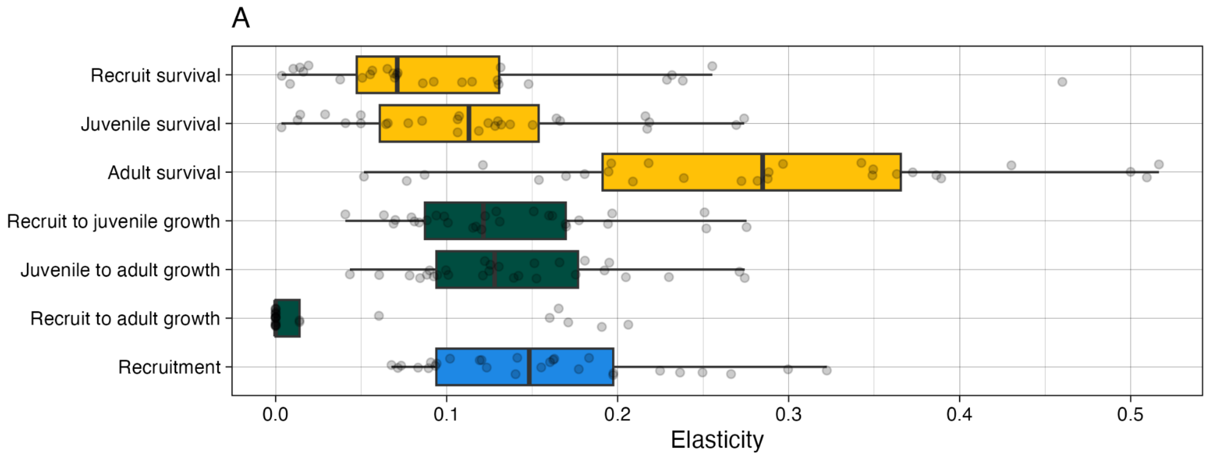


**Figure S4**. Elasticities. Impact of vital rate changes on population growth rate, expressed as an elasticity of each specified rate across all 28 models.

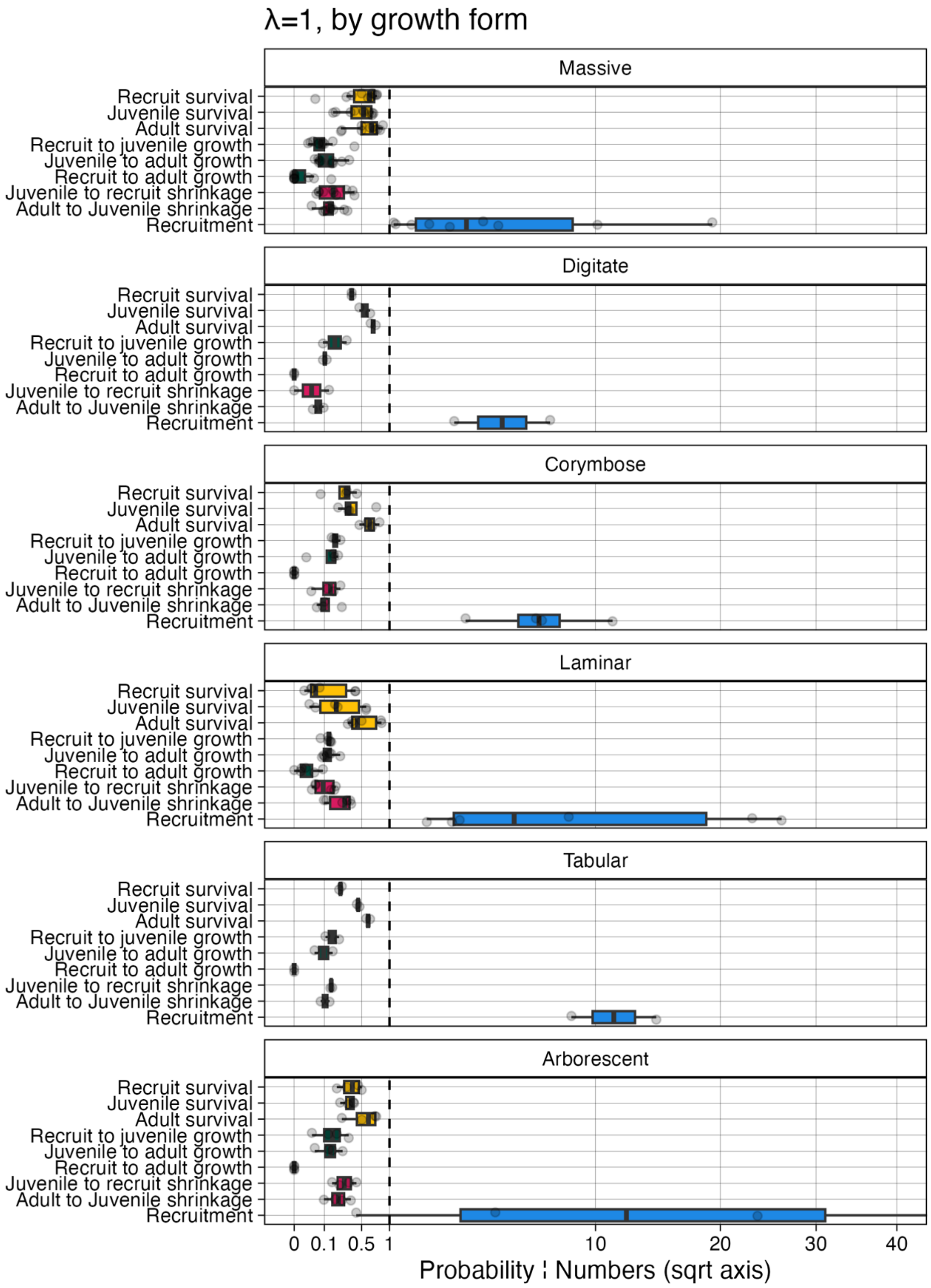


**Figure S5**. Boxplots for transition cells for the 28 species matrices summarized by growth form. Color coding matches Fig. 1. Points are for each species.

**Supporting Results.** A key uncertainty with the matrices used in this study was the assumption of initial population stasis (λ = 1). Increasing population growth rate by 5% when not at stasis may require different amounts of structural and supplemental change. We therefore also estimated recruitment for each population matrix required for λ levels ranging from decline, through stasis, to growth (i.e., λ = 0.9, 0.95, 1, 1.05, 1.1). We then asked what was required to increase population growth by 5%, which equates to increasing λ to 0.945, 0.9975, 1.05, 1.1025, 1.155, respectively. Initial λ did not have a large effect on the magnitude of the structural and supplemental approaches (Fig. S6). However, there were some general trends. For example, structural multipliers for growth and recruitment needed to be marginally greater when populations were more in decline; and more adults would need to be outplanted, but fewer recruits and juveniles when populations were more in decline.

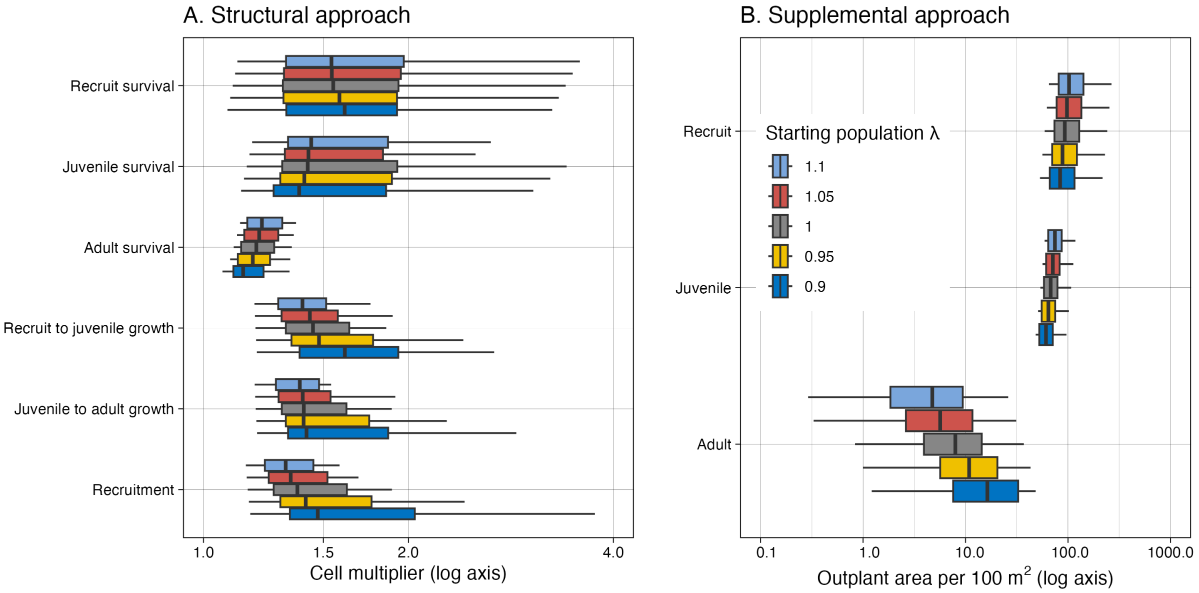


**Figure S6**. Boxplots for (A) transition cells and (B) outplant area required to increase population growth by 5% for the 28 species matrices summarized by starting population growth rate.
